# scACORN: Context-engineered agent orchestration of specialized small language models for single-cell transcriptomic interpretation

**DOI:** 10.64898/2026.09.10.750801

**Authors:** Arash Rasti-Meymandi, Sepideh Nahali, Eustache Paramithiotis, Angela M. Cheung, Elham Dolatabadi

**Affiliations:** Department of Medicine, University of Toronto, Toronto, Ontario, Canada; School of Health Policy and Management, Faculty of Health, York University, Toronto, Ontario, Canada; York University, Toronto, Ontario, Canada; CellCarta, Montreal, Quebec, Canada; Vector Institute, Toronto, Ontario, Canada

**Keywords:** single-cell transcriptomics, language models, agentic systems, small language models, contrastive learning

## Abstract

Single-cell atlases now exceed 66 million cells, but turning a ranked expression profile and a free-form biological question into a reliable, evidence-grounded answer remains unsolved. Scaling a single model does not resolve this, because single-cell interpretation is a heterogeneous family of tasks whose correct answer depends on tissue, cohort, perturbation and annotation resolution. Here we present scACORN, an agentic alternative to monolithic single-cell language models that combines specialized small language models with context-engineered agent orchestration for their selection and composition at inference time. Each expert is built in two stages: domain-aligned contrastive adaptation fits a pretrained cell-to-text backbone to the transcriptomic geometry of a target dataset, and geometry-preserving specialization learns question-conditioned biological completions without eroding that geometry. A fixed orchestrating language model agent then selects and combines experts under a natural-language playbook that is itself optimized from textual feedback, with no gradient updates to the orchestrator. Across 10 Tabula Sapiens tissues, domain alignment raised transfer macro-F1 from 0.36 to 0.64 and Recall@5 from 0.87 to 0.97; specialized experts reached 0.89 mean exact-match annotation accuracy; and playbook optimization reduced unsupported gene citations from 14.5% to 3.5%. Our findings support specialization and orchestration as complementary responses to the heterogeneity and evidentiary demands of single-cell analysis.

## 1 Introduction

Single-cell atlases have made cellular heterogeneity measurable at scale (Regev et al. 2017; Rood et al. 2025; Tabula Sapiens Consortium 2022). Between 2020 and 2025 the Human Cell Atlas grew more than elevenfold, from 5.8 million to over 66 million cells and beyond 300 TB (Human Cell Atlas Data Portal 2020; Human Cell Atlas 2025). These resources have enabled detailed characterization of the cellular identities, states and responses that shape tumor heterogeneity, immunotherapy response and human development (Tirosh et al. 2016; Sade-Feldman et al. 2018; Cao et al. 2020). Yet atlas-scale analysis requires more than assigning a label to an individual expression profile. It requires resolving closely related lineages, interpreting transcriptional programs in the context of tissue, disease and perturbation, comparing cells across experimental conditions, and linking biological conclusions to the genes that support them. As single-cell datasets increase in both scale and biological diversity (Regev et al. 2017; Tabula Sapiens Consortium 2022), the principal computational challenge is shifting from data generation and representation toward reproducible, context-sensitive and evidence-grounded interpretation (Svensson et al. 2018; Lähnemann et al. 2020).

Foundation models offer a potential route to scale this interpretation. Geneformer (Theodoris et al. 2023), scGPT (Cui et al. 2024) and scFoundation (Hao et al. 2024) learn general-purpose transcriptomic representations, whereas Cell2Sentence and Cell-o1 connect single-cell profiles to natural-language generation and reasoning (Levine et al. 2024; Fang et al. 2025). However, increasing model scale does not, by itself, resolve the interpretation problem. Zero-shot transfer remains weak, perturbation prediction does not beat linear baselines, and task-specific models frequently match or exceed far larger ones (Kedzierska et al. 2025; Ahlmann-Eltze et al. 2025). Single-cell interpretation comprises a heterogeneous set of tasks, and a single biological question may require several forms of expertise. The significance of an expression program may vary with tissue, cohort, modality, perturbation and annotation resolution (Luecken et al. 2022; Domínguez Conde et al. 2022; Abdelaal et al. 2019; Hrovatin et al. 2025). Monolithic adaptation therefore conflates two requirements: acquiring precise local expertise and determining which expertise applies to each profile and question.

Here we formulate single-cell interpretation as the *agentic composition of specialized* biological expertise. Given one or more transcriptomic profiles represented by ranked gene lists and a free-form biological question, the system must identify the relevant biological subproblems, assign each profile to the appropriate specialists, and synthesize a coherent answer in which molecular claims remain traceable to the corresponding measurements. We introduce scACORN (**s**ingle-**c**ell **A**gentic **C**ontext **OR**chestration **N**etwork), an agentic specialist–orchestrator designed to meet these requirements (Fig. 1). scACORN constructs reusable cell-to-text experts from target-domain data and selects and composes them at the level of individual profiles. In contrast to conventional tool-using agents, which operate over a predefined collection of fixed tools (Yao et al. 2023; Schick et al. 2023), scACORN builds a repertoire of specialized biological experts and dynamically orchestrates them according to the biological context and question at hand.

**Fig. 1.**
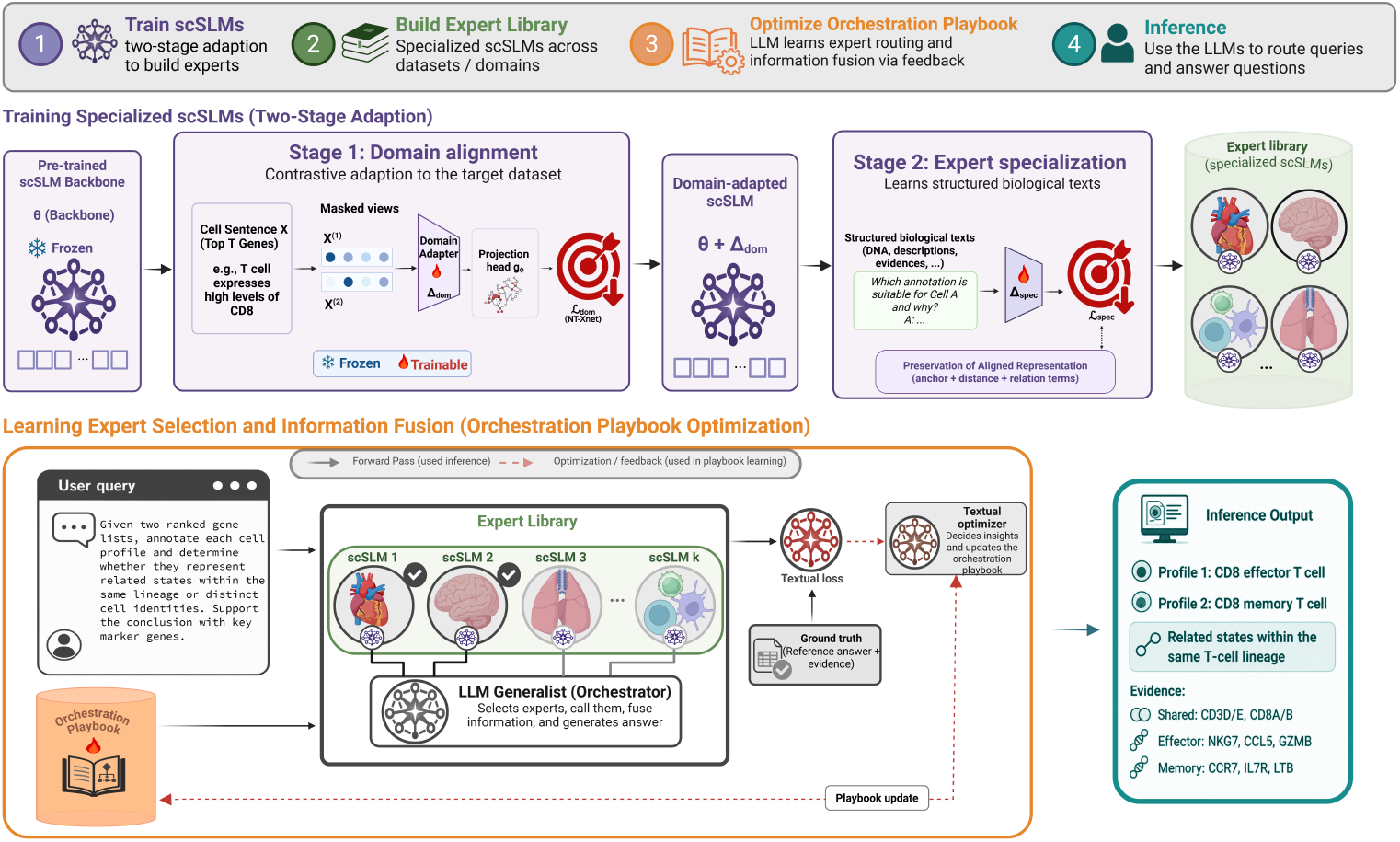
Overview of scACORN (single-cell Agentic Context ORchestration Network). **1**, Domain-aligned contrastive adaptation fits a pretrained cell-to-text backbone to a target transcriptomic domain, and geometry-preserving specialization converts it into a question-conditioned expert. **2**, The trained experts form the collection available to the orchestrator. **3**, The orchestration playbook is optimized from textual feedback while the expert and orchestrator parameters remain fixed. **4**, At inference, the orchestrator selects one or more experts and synthesizes their outputs into a free-form answer with profile-linked molecular evidence. Solid arrows indicate inference flow, and dashed arrows indicate feedback used for playbook optimization.

scACORN separates the acquisition of local biological expertise from the policy used to select and integrate that expertise. Experts are built from a pretrained single-cell small language model (scSLM) backbone in two stages. Domain-aligned contrastive adaptation fits a pretrained cell-to-text backbone to the transcriptomic geometry of a target dataset, establishing competence within a defined domain. Geometry-preserving specialization then converts that competence into question-conditioned predictions and gene-level rationales while explicitly penalizing drift from the aligned representation. The resulting tissue-, dataset-, modality- or task-specific models form the set of experts available to the orchestrator. Composition is performed by one of two fixed language-model orchestrators, GPT-5.4 mini or Claude Sonnet 4.5, guided by an explicit natural-language playbook that is optimized from structured textual feedback rather than gradients (Zhang et al. 2025; OpenAI 2026; Anthropic 2025). Because experts are constructed independently and share a textual interface, the framework provides a modular basis for incorporating additional specialists.

In total we trained 20 adapters spanning 10 Tabula Sapiens tissues (Tabula Sapiens Consortium 2022), generated and scored approximately 170,000 held-out cell annotations against 12 baseline systems, and evaluated composition on 80 held-out free-form questions under two orchestrating language models. Domain alignment increased mean cell-type classification macro-F1 from 0.356 to 0.643 and Recall@5 from 0.868 to 0.968, bringing a language-model representation into the range of purpose-built ranked-gene encodings. Geometry-preserving specialization produced tissue experts with 0.90 pooled exact cell-type accuracy and an evidence-gene F1 of 0.82, while retaining hidden-space Recall@10 at 0.969 at the reported precision. On held-out free-form questions, including 42 that required two or three experts, playbook optimization reduced unsupported support-gene citations from 14.5% to 3.5% and increased ontology-aware agreement from 0.754 to 0.843. scACORN achieved the highest ontology agreement, evidence alignment and nuanced-answer quality among the evaluated general-purpose and transcriptomic models.

## 2 Results

We evaluated scACORN (Fig. 1) across the three linked stages of its workflow; *First*, we asked whether contrastive domain alignment could adapt a pretrained cell-to-text backbone to the transcriptomic structure of a target tissue. *Second*, we tested whether the aligned backbone could be converted into a cell-level reasoning expert that produces accurate labels and grounded marker rationales. *Third*, we evaluated whether textual orchestration could answer free-form questions by selecting and combining the appropriate experts from a library of specialized models.

### 2.1 Experimental Design

All experiments used Tabula Sapiens as a common biological substrate (Tabula Sapiens Consortium 2022). The domain-alignment benchmark included cells from 10 tissues. To reduce leakage from experimentally linked observations, cells belonging to the same acquisition group, including cells from the same 10x run, were assigned to the same training, validation or test partition. The expert-specialization benchmark evaluated tissue-specific biological completions on held-out cells from the same 10-tissue collection. External model comparisons used matched held-out profiles within each tissue so that all methods received the same ranked-gene inputs.

The orchestration benchmark comprised 80 fixed held-out questions: 38 single-expert questions and 42 compositional questions. Single-expert questions required one tissue-specialized expert, whereas compositional questions required the selection and combination of two or three experts. The reference expert set associated with each question was used only by the evaluator for routing diagnostics. All compared systems received the same profiles, questions and evaluation rubric.

To test whether orchestration performance depended on a particular model scale or provider, we evaluated `GPT-5.4 mini` as a compact OpenAI orchestrator and `Claude Sonnet 4.5`as a larger-capacity Anthropic orchestrator (OpenAI 2026; Anthropic 2025). The set of experts, held-out questions and evaluation procedure were fixed across the two configurations.

For domain alignment, we compared Rank PCA, TF-IDF followed by truncated SVD, the unadapted pretrained cell-to-text backbone, the hidden representations produced after domain alignment, and the corresponding learned projection space. For expert specialization, we compared scACORN experts with open-source instruction-tuned models from the Mistral, Qwen, Gemma and Llama families (Jiang et al. 2023; Qwen Team 2025; Yang et al. 2025; Gemma Team 2024; Dubey et al. 2024), closed-source models (OpenAI 2024, 2025), and specialized transcriptomic language models (Fang et al. 2025; Rizvi et al. 2025). Primary metrics were aligned with the claim tested by each component: transfer macro-F1 and Recall@5 for domain alignment; exact label agreement and evidence grounding for expert specialization; and ontology-aware agreement, evidence alignment, marker coverage and nuanced answer quality for orchestration. Paired model comparisons used sample-level randomization tests against the strongest non-scACORN baseline. Formal metric definitions are provided in Appendix C.

### 2.2 Domain Alignment Fits A Language-Model Backbone to A Target Single-Cell Domain

Because expert specialization operates on the adapted LLM state rather than on TF-IDF, PCA or another fixed feature space, we evaluated domain alignment as an upstream component of expert construction. Specifically, we asked whether contrastive adaptation (Fig. 2) improved the local structure of the unfitted cell-to-text representation and brought it into the range of conventional ranked-gene representations before question-conditioned specialization.

**Fig. 2.**
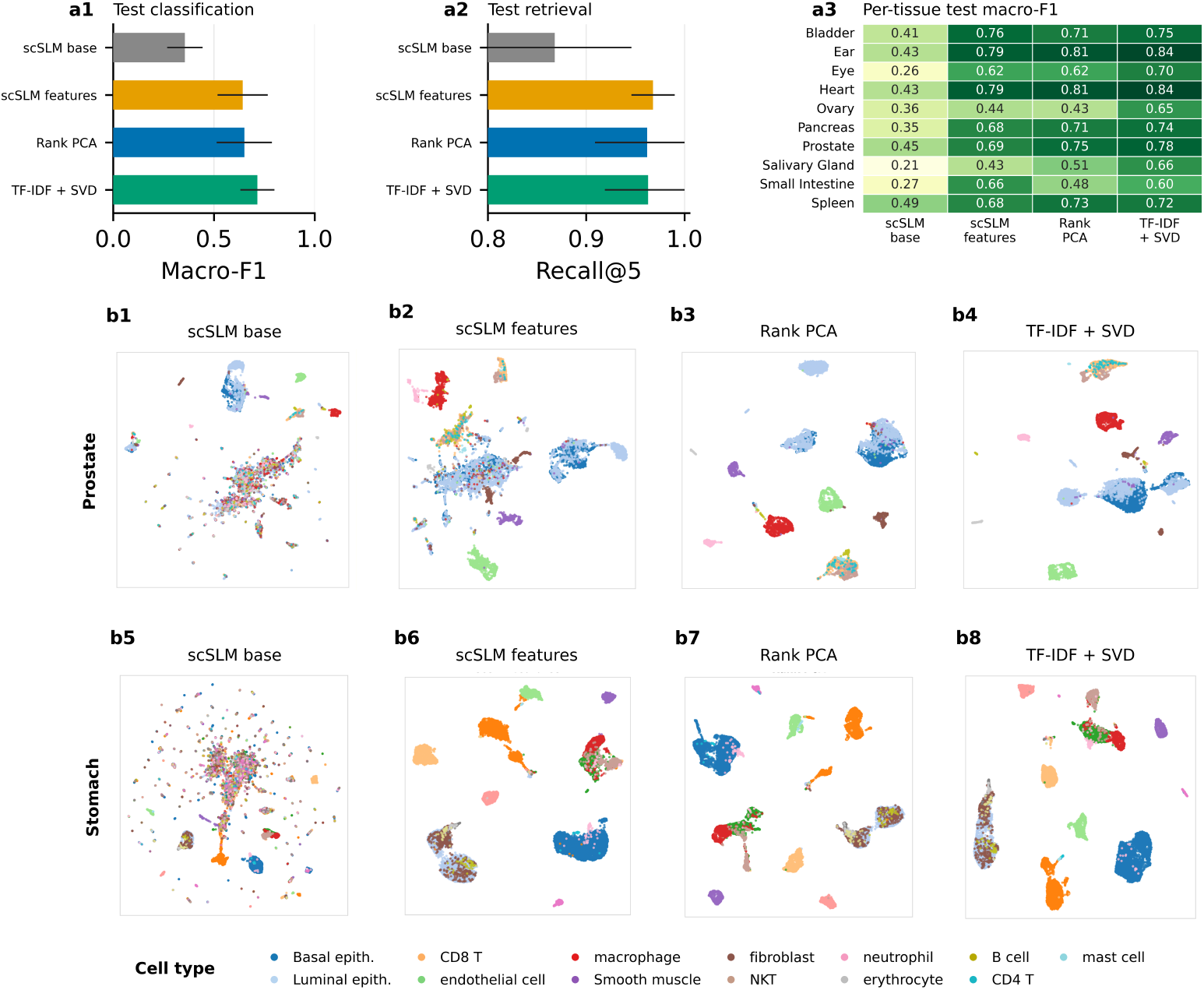
Domain alignment improves cell-type structure in cell-to-text representations. **a1**, Test classification macro-F1 for the unadapted single-cell small language model (scSLM) backbone, domain-aligned scSLM hidden features, Rank PCA and TF-IDF + SVD representations. **a2**, Test retrieval Recall@5 for the same four representations. **a3**, Per-tissue test macro-F1 across the 10 evaluated tissues. Bars in **a1** and **a2** show the mean across tissues; error bars indicate the s.d. across tissues. **b1–b4**, UMAP projections (McInnes et al. 2018) of prostate cells represented by the unadapted scSLM backbone, domain-aligned hidden features, Rank PCA and TF-IDF + SVD, respectively. **b5– b8**, Corresponding projections for stomach cells. Each point represents one cell, and colors denote reference cell-type annotations.

Across the 10 tissues, the unfitted scSLM backbone obtained a mean classification macro-F1 of 0.356± 0.088. Domain alignment increased macro-F1 to 0.643 ±0.125 for the adapted hidden features and 0.640±0.171 for the learned projection space. TF-IDF + SVD reached 0.716 ±0.085, and Rank PCA reached 0.651 ±0.137 (Fig. 2(a1,a3)). Thus, contrastive adaptation substantially improved the pretrained language-model representation and produced classification structure comparable to the conventional representations, while retaining a state that could be reused by the downstream SLM expert.

The retrieval analysis showed a similar pattern (Fig. 2(a2)). Adapted hidden features achieved the highest mean Recall@5, 0.968 ±0.022, compared with 0.868 ±0.078 for the unfitted backbone, 0.963 ± 0.045 for TF-IDF + SVD, 0.962 ± 0.053 for Rank PCA and 0.951 ± 0.052 for the learned projection space. These results do not establish domain alignment as a new general-purpose embedding method; rather, they show that it fits the cell-to-text backbone to the target biological domain while retaining competitive local separation.

Representative prostate and stomach embeddings provided a qualitative view of the same effect (Fig. 2(b1–b8)). The unfitted scSLM representation showed weaker cell-type organization, whereas the adapted hidden features formed more coherent local neighborhoods.

### 2.3 Expert Specialization Improves cell-type Reasoning and Molecular Grounding

We next asked whether the domain-aligned backbone could be converted into a question-conditioned biological expert. Across the 10-tissue expert-specialization benchmark, scACORN experts achieved a mean exact-match rate of 0.893 ± 0.096. Performance varied across tissues, ranging from 0.990 in bladder to 0.719 in small intestine and 0.760 in salivary gland. Detailed tissue-level results are reported in Appendix A.

scACORN also outperformed substantially larger general-purpose and transcriptomics-oriented models on label quality. Across the full evaluation, scA-CORN reached 0.95 canonical accuracy, 0.90 exact accuracy, 0.95 ontology credit, 0.73 macro-F1 and 0.96 weighted F1. The ontology-credit and macro-F1 comparisons are summarized in Fig. 3(a,b). The strongest open-source baseline, `Mistral-Small-3.2-24B-Instruct-2506`, reached 0.76 canonical accuracy and 0.41 macro-F1. The strongest specialized baseline, Cell-o1, reached 0.70 canonical accuracy and 0.41 macro-F1, whereas the best closed-source result reached 0.54 canonical accuracy and 0.59 ontology credit. The larger difference in macro-F1 is consistent with improved performance beyond the dominant cell classes.

**Fig. 3.**
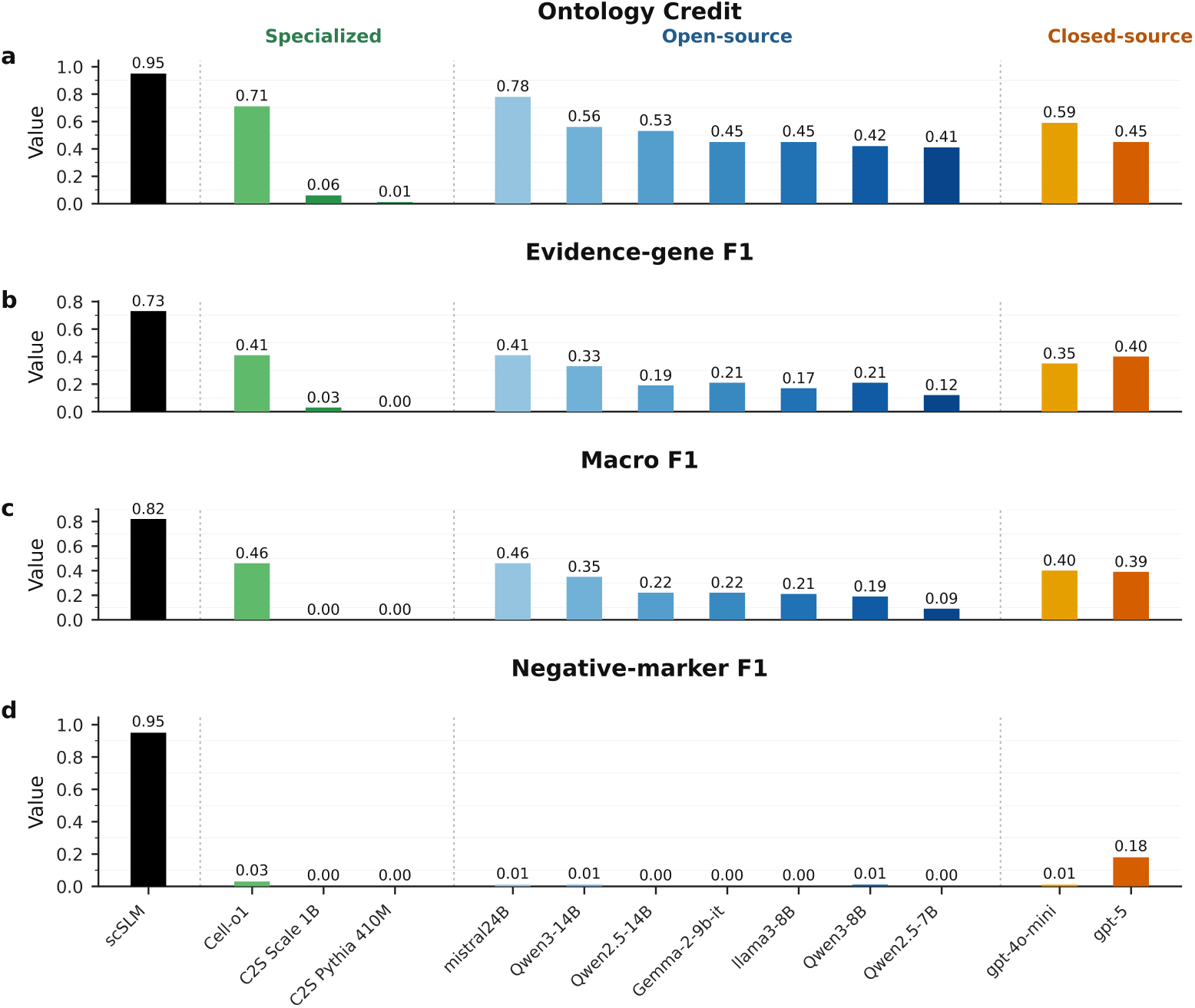
Expert specialization improves cell-type reasoning and molecular grounding. **a–d**, Performance of the scACORN’s experts (scSLMs) compared with specialized transcriptomic models, open-source instruction-tuned language models and closed-source language models on matched heldout cell profiles. **a**, Ontology credit, which awards partial credit to predictions that are biologically related to the reference cell type. **b**, Evidence-gene F1, measuring agreement between genes cited in the generated rationale and the reference supporting-gene set. **c**, Macro-F1 across cell-type classes. **d**, Negative-marker F1, measuring agreement between predicted and reference genes used to exclude competing cell identities. Higher values are better; values are shown above the bars.

The advantage of expert specialization extended to the molecular evidence in the generated answers (Fig. 3 (c,d)). scACORN obtained evidence-gene precision, recall and F1 of 0.82, 0.82 and 0.82, respectively. The strongest open-source baseline reached an evidence-gene F1 of 0.46, the strongest closed-source baseline reached 0.40, and Cell-o1 reached 0.46. scACORN also obtained negative-marker precision, recall and F1 of 0.96, 0.94 and 0.95, respectively.

Qualitative examples further illustrated the distinction between label prediction and grounded reasoning. scACORN identified bladder urothelial cells using epithelial programs including *KRT19, CLDN4* and *TACSTD2*; fibroblasts using *CFD, DCN* and *MGP*; and heart endothelial cells using *PECAM1, VWF* and *ENG*. In a multi-profile question, the model separated an ileal enterocyte profile supported by *APOA1, APOA4, SLC5A1* and *FABP2* from a pericyte profile supported by *RGS5, NOTCH3* and *CALD1*. Full qualitative interactions are provided in Appendix D.

### 2.4 Textual Orchestration Improves Free-Form Biological Question Answering

Finally, we evaluated whether a language-model orchestrator guided by an optimized textual playbook could select and combine specialized experts to answer free-form biological questions. The held-out benchmark included both questions requiring one expert and compositional questions requiring two or three expert calls. The latter setting requires the system to determine which domain-specific knowledge is relevant, associate each profile with the appropriate expert and synthesize evidence across multiple expert outputs.

As indicated in Fig. 4, playbook optimization improved the orchestrator’s use of expert evidence and the compatibility of its predictions with the reference labels (Fig. 4). For GPT-5.4 mini, the input-grounded support-gene score increased from 0.855 with the initial playbook to 0.965 at the prespecified final endpoint, reducing the unsupported support-gene citation rate from 0.145 to 0.035. Ontology-aware agreement increased from 0.754 to 0.843. Claude Sonnet 4.5 showed a similar overall improvement. Although performance varied across intermediate versions of the playbook, the final version outperformed the initial version on both measures.

**Fig. 4.**
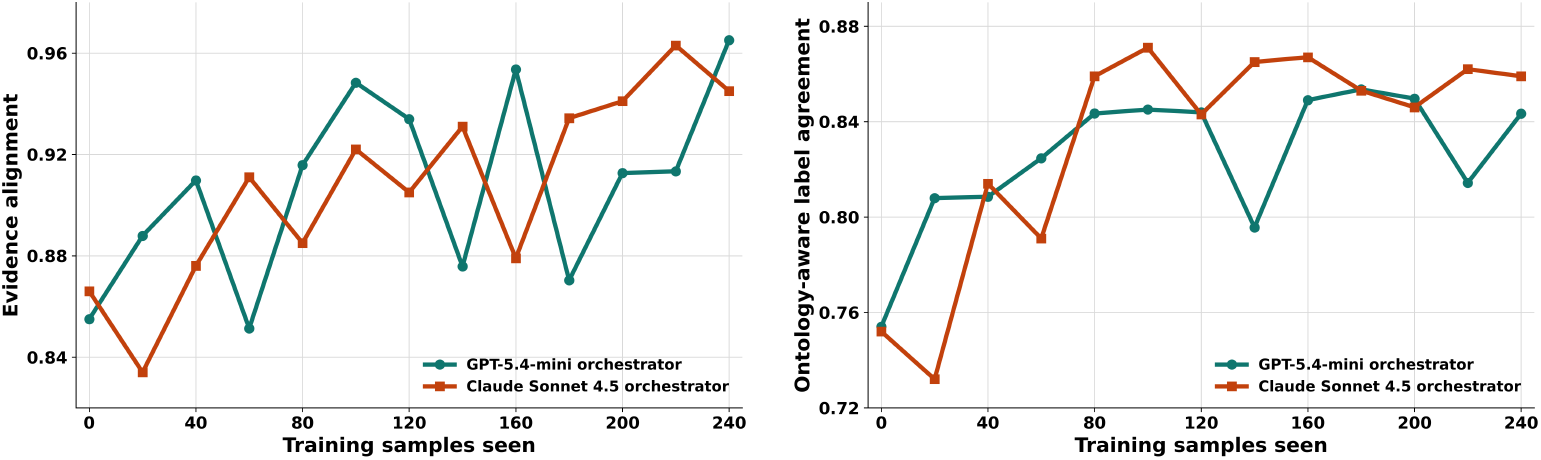
Textual playbook optimization improves evidence grounding and ontology-aware agreement under two orchestrators. Left, input-grounded support-gene score at successive playbook-optimization checkpoints. The unsupported support-gene citation rate is one minus this score. Right, ontology-aware label agreement at the same checkpoints. Curves show results for the GPT-5.4-mini and Claude Sonnet 4.5 orchestrators. Dashed horizontal lines indicate performance with the corresponding initial playbook. Each checkpoint was evaluated on the same fixed held-out set; the final endpoint was specified in advance, and intermediate checkpoints were not used to select the reported playbook.

Across all 80 held-out questions, Claude-orchestrated scACORN achieved an ontology score of 0.850, an evidence-alignment score of 0.603, a marker-coverage score of 0.814 and a nuanced-answer score of 0.748. The corresponding GPT-5.4-mini results were 0.843, 0.567, 0.798 and 0.744. Claude Sonnet 4.5 therefore produced the strongest ontology, evidence-alignment and nuanced-answer results among the evaluated systems. GPT-5, the strongest standalone closed-source baseline, obtained 0.707, 0.328, 0.834 and 0.504 and retained the highest aggregate marker coverage. Cell-o1 obtained 0.457, 0.283, 0.550 and 0.480.

Even with an empty playbook, the initial prompt sometimes led the orchestrator to select an appropriate expert. Playbook optimization made this selection more reliable, which helps explain the large improvement in evidence alignment for compositional questions: once the appropriate experts were selected, their outputs were already strongly aligned with the reference evidence. The other metrics changed more moderately, and marker coverage was less consistent.

For single-expert questions (*n* = 38), the optimized Claude-orchestrated scACORN achieved the highest score on all four metrics. For compositional questions (*n* = 42), it achieved the highest ontology, evidence-alignment and nuanced-answer scores, whereas the empty-playbook Claude condition had the highest marker coverage (Table 1).

**Table 1.** Textual orchestration performance across orchestrators, question types and playbook conditions. **a**, Single-expert questions (*n* = 38), each requiring one tissue-specialized expert. **b**, Compositional questions (*n* = 42), each requiring two or three experts. For scACORN, the model in parentheses denotes the orchestrating language model. Values are mean scores across held-out questions; higher is better. The best result for each metric within each panel is shown in bold. Empty indicates that no playbook instructions were supplied; optimized denotes the final optimized playbook. Evidence align., weighted set-F1 agreement between predicted and reference supporting and negative-marker sets; Marker cov., weighted recall of reference supporting and negative markers mentioned in the generated answer.

| Category | Model | Ontology ↑ | Evidence align. ↑ | Marker cov. ↑ | Nuanced ↑ |
| --- | --- | --- | --- | --- | --- |
| **a. Single-expert questions** |  |  |  |  |  |
| *Open-source* | Mistral-Small-3.2-24B-Instruct | 0.593 | 0.311 | 0.506 | 0.435 |
|  | Qwen3-14B | 0.460 | 0.200 | 0.342 | 0.371 |
|  | Llama-3.1-8B-Instruct | 0.216 | 0.135 | 0.296 | 0.273 |
| *Closed-source* | GPT-4o-mini | 0.480 | 0.308 | 0.388 | 0.400 |
|  | GPT-5 | 0.633 | 0.290 | 0.865 | 0.486 |
| *Specialized* | Cell-o1 | 0.411 | 0.259 | 0.556 | 0.476 |
|  | scACORN (GPT-5.4-mini; empty) | 0.780 | 0.537 | 0.902 | 0.699 |
|  | **scACORN (GPT-5.4-mini; optimized)** | 0.832 | 0.632 | 0.904 | 0.747 |
|  | scACORN (Claude Sonnet 4.5; empty) | 0.751 | 0.543 | 0.914 | 0.709 |
|  | **scACORN (Claude Sonnet 4.5; optimized)** | **0.841** | **0.678** | **0.922** | **0.753** |
| **b. Compositional questions** |  |  |  |  |  |
| *Open-source* | Mistral-Small-3.2-24B-Instruct | 0.491 | 0.301 | 0.461 | 0.377 |
|  | Qwen3-14B | 0.569 | 0.193 | 0.375 | 0.381 |
|  | Llama-3.1-8B-Instruct | 0.338 | 0.113 | 0.209 | 0.280 |
| *Closed-source* | GPT-4o-mini | 0.543 | 0.257 | 0.334 | 0.384 |
|  | GPT-5 | 0.773 | 0.364 | 0.807 | 0.520 |
| *Specialized* | Cell-o1 | 0.499 | 0.305 | 0.544 | 0.482 |
|  | scACORN (GPT-5.4-mini; empty) | 0.614 | 0.019 | 0.793 | 0.513 |
|  | **scACORN (GPT-5.4-mini; optimized)** | 0.853 | 0.508 | 0.702 | 0.741 |
|  | scACORN (Claude Sonnet 4.5; empty) | 0.622 | 0.139 | **0.808** | 0.553 |
|  | **scACORN (Claude Sonnet 4.5; optimized)** | **0.858** | **0.535** | 0.716 | **0.744** |

With optimized playbooks, Claude Sonnet 4.5 outperformed GPT-5.4 mini on all four metrics in both subsets.

## 3 Discussion

We introduce scACORN, which constructs specialized scSLMs and uses a context-engineered agent to orchestrate their selection and composition for single-cell transcriptomic interpretation. Each expert is trained within a defined biological domain, while the orchestrator selects and combines those relevant to the profiles and question at hand. The shared textual interface preserves the provenance of molecular evidence by linking each contribution to its source profile and expert. The resulting design supports broader biological questions through the composition of focused expertise. This modularity also allows the system to accommodate new experts without altering its overall structure. For example, a new scSLM can be constructed by applying domain alignment and expert specialization to a new tissue, dataset or task, then added to the available expert set together with a textual description of its scope and input requirements. Because the playbook specifies how and when experts should be invoked, the orchestrator can consider the new specialist at inference, although routing reliability as the expert set expands remains to be evaluated.

Single-cell interpretation requires local adaptation without erasing the transcriptomic structure that distinguishes related cell populations (Hrovatin et al. 2025; Li and Hoiem 2016). Our findings support this rationale. Domain alignment organized the language-model backbone around tissue-specific transcriptomic structure, providing a biologically appropriate starting point for expert specialization. The preservation objective then allowed experts to learn question-conditioned outputs without measurable loss of this structure under the present evaluation. This was particularly important for distinguishing closely related populations, whose identities depend on both expressed markers and the absence of competing lineage programs (Abdelaal et al. 2019; Domínguez Conde et al. 2022). Accordingly, the compact experts improved not only label quality but also the recovery of supporting and exclusionary evidence relative to substantially larger models. At the system level, playbook optimization primarily improved how this evidence was used, substantially reducing unsupported gene citations across both single- and multi-expert questions.

This study provides an initial validation of scACORN in a controlled, multi-tissue setting and identifies several priorities for further evaluation. Deriving all experts from Tabula Sapiens enabled matched comparisons across tissues and system components; validation in independent atlases, disease and perturbation cohorts, and data generated using different protocols will now be important for establishing robustness to dataset shift. Similarly, our benchmarks isolate cell identification and evidence synthesis, providing a foundation for prospective studies of whether the framework can support biological discovery and generate hypotheses that withstand expert review and experimental validation. The current set of experts also offers a tractable testbed for orchestration, whereas larger collections will introduce challenges related to overlapping expertise, conflicting outputs and inference cost. Our trace-based routing controls indicate non-random expert selection, but complete generative reruns under oracle, random and disabled routing will be needed to determine its causal contribution to answer quality. Finally, gene-level grounding provides a transparent measure of whether cited evidence is present in the input or reference set, but should not be interpreted as evidence of biological specificity, mechanism or causality.

Future work should evaluate broader sets of experts across independent datasets, disease and perturbation cohorts, rare-cell populations and multimodal measurements such as CITE-seq, spatial transcriptomics and chromatin accessibility. As the number of experts grows, orchestration will need to account explicitly for expert overlap, disagreement, uncertainty and computational cost. This reframes biological model scaling as a problem of organizing and coordinating expertise rather than continually enlarging a single model.

## 4 Methods

### 4.1 Problem Formulation and Framework Overview

We consider grounded cell-level biological question answering. The index *i* denotes an example or cell. Each example contains a ranked single-cell transcriptomic profile *x*_*i*_, a natural-language question *q*_*i*_, optional task context *c*_*i*_, and, during supervised training, a target textual answer *y*_*i*_. Given one or more profiles and a question, the objective is to generate a biologically compatible answer whose supporting evidence is traceable to the corresponding observed profile. The system must also identify the relevant domain-specific expert or combination of experts and avoid citing genes that are unsupported by the associated ranked input. The target answer may contain a cell-type label, supporting genes, negative markers, tissue context, alternative interpretations, or a concise biological rationale.

scACORN separates representation adaptation, expert construction, and run-time reasoning into three sequential components. Domain alignment first adapts a pretrained cell-to-text backbone to the transcriptomic structure of a target dataset or tissue. Geometry-preserving expert specialization then learns question-conditioned biological completions without erasing the aligned representation. Finally, textual expert orchestration registers the resulting specialists as tools and selects one or more of them through an explicitly optimized playbook.

Unless stated otherwise, *r* indexes gene rank, *m* indexes target tokens, *v* indexes augmented views, *k* indexes experts, *p* indexes input profiles, *j* indexes tool calls, and *t* indexes textual-optimization iterations. The same expert-construction procedure is applied independently to each domain-specific expert; we omit the expert identifier during domain alignment and specialization for clarity.

### 4.2 Cell-to-Text Representation

Each cell is represented by its top *T* expressed genes,

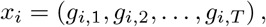

where *g*_*i,r*_ is the gene at rank *r* for cell *i*, and genes are ordered by decreasing expression. We use *T* = 200 in all experiments. A deterministic prompt template *T* converts the ranked profile, question, and optional context into a textual prompt:

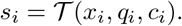

During domain alignment, no question or task context is supplied, so the corresponding prompt is *T* (*x*_*i*_, ∅, ∅).

Let *θ* denote the frozen parameters of the pretrained causal language-model backbone, and let Δ denote an active adapter or ordered collection of adapters. For a prompt *s*, we write **h**_*θ*,Δ_(*s*) ∈ ℝ^*d*^ for the hidden state of its final non-padding token, where *d* is the backbone hidden dimension. The full sequence-level definition is provided in Appendix B.1. This formulation follows the cell-to-text paradigm and supports parameter-efficient domain alignment and expert specialization (Levine et al. 2024; Rizvi et al. 2025).

### 4.3 Domain-Aligned Contrastive Adaptation

Domain alignment adjusts the pretrained backbone to the local transcriptomic structure of a target domain before supervised reasoning is introduced. Let *A* denote the stochastic augmentation operator applied to ranked-gene profiles. For each cell *x*_*i*_, we draw two independent views,

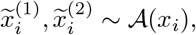

where A applies biologically conservative perturbations, including gene dropout, local rank swaps, and random truncation or subsampling.

The augmented profiles are converted into cell-to-text prompts and encoded using a trainable domain-alignment LoRA adapter Δ_dom_ (Hu et al. 2022). A projection head 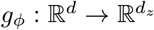, parameterized by *ϕ*, maps each prompt representation to a *d*_*z*_-dimensional contrastive space. For any nonzero vector **u**, define norm(**u**) = **u***/*∥**u**∥_2_, where ∥ · ∥_2_ is the Euclidean norm. The normalized embedding for view *v* ∈ {1, 2} is

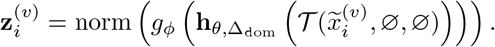

For a mini-batch of *N* cells, we optimize the NT-Xent objective (Chen et al. 2020):

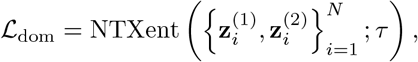

where *N* is the number of cells in the contrastive mini-batch and *τ >* 0 is the temperature. The two views of the same cell form a positive pair, whereas all other views in the batch act as negatives. The expanded objective is given in Appendix B.2.

Only Δ_dom_ and *ϕ* are optimized. In our implementation, LoRA modules are applied to the attention projections in the upper half of the transformer layers. The projection head is used only during domain-alignment training and representation evaluation.

Checkpoints are selected using validation Recall@10 in the projection space under cosine similarity. This criterion reflects the purpose of domain alignment: adapting the language-model representation to the target domain rather than directly optimizing cell-type prediction.

### 4.4 Geometry-Preserving Expert Specialization

Expert specialization converts the domain-aligned backbone into a question-conditioned biological reasoning expert. The domain-alignment adapter Δ_dom_ remains active and frozen, while a second adapter Δ_spec_ is trained for the target completion tasks.

For training example *i*, the target answer is tokenized as

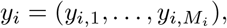

where *M*_*i*_ is the number of target tokens and *y*_*i,<m*_ denotes the prefix preceding token *m*. For a specialization mini-batch *ℬ*_spec_, we optimize the completion-only causal language-modeling loss

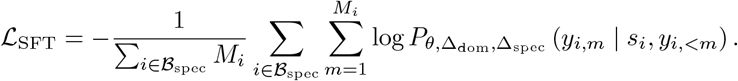

Here, *P*_*θ*,Δdom_,Δ_spec_ denotes the model next-token distribution. Prompt tokens are excluded from the loss. Training targets contain the final cell label and, when available, supporting genes, negative markers, tissue information, confidence, and a short rationale.

### 4.5 Representation-Preservation Objective

Expert specialization can improve completion quality while distorting the domain geometry learned through domain alignment, reflecting the broader problem of representation drift during sequential adaptation (Li and Hoiem 2016). We therefore compare the normalized prompt representation of the current specialized expert with that of the frozen domain-aligned reference model. The three preservation terms draw on pointwise feature alignment, moment matching, and relational knowledge preservation (Sun and Saenko 2016; Park et al. 2019).

We preserve the domain-aligned representation at three complementary levels:

1. **pointwise alignment**, which preserves each prompt representation;
2. **distribution alignment**, which preserves batch means and componentwise variances; and
3. **relational alignment**, which preserves pairwise similarities among prompts in the same mini-batch.

The complete specialization objective is

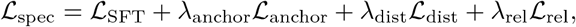

where *λ*_anchor_, *λ*_dist_, *λ*_re_ ≥ 0 weight the pointwise, distributional, and relational preservation losses, respectively. Their exact definitions are provided in Appendix B.3. Only Δ_spec_ is updated during expert specialization. Checkpoints are selected using validation completion loss, while hidden-space retrieval on a held-out domain split is monitored as an auxiliary measure of representation retention.

### 4.6 Textual Expert Orchestration

After expert specialization, each specialized model is made available to the orchestrator as a textual tool, following the broader tool-augmented agent paradigm in which an LLM selects and invokes external capabilities during reasoning (Yao et al. 2023; Schick et al. 2023).

Let

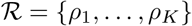

denote the set of *K* experts, where descriptor *ρ*_*k*_ specifies expert *k*’s dataset or tissue, supported tasks, adapter location, and routing description. A query may contain *P* ranked-gene profiles,

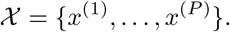

Given a free-form question *q*, profile set, *X* optional context *c*, expert set, *ℛ* and textual playbook, *P* the selected fixed orchestrating LLM defines a playbook-conditioned textual policy Π:

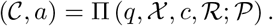

Here,

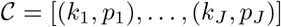

is the ordered tool trace containing *J* expert calls, *k*_*j*_∈ {1, …, *K*} is the expert selected at call *j, p*_*j*_ ∈ {1, …, *P*} is the profile supplied to that expert, and *a* is the synthesized final answer.

We instantiated this policy separately with GPT-5.4 mini and Claude Sonnet 4.5. In both configurations, the orchestrator parameters remained fixed and only the textual playbook was updated. Both configurations used the same experts and tool interface. The policy receives descriptions of all available experts but does not receive the identity of the correct expert. It must therefore determine which expert or combination of experts is appropriate for the query. This component is the mechanism that enables accurate responses to free-form questions: instead of relying on a single general model for all biological contexts, the orchestrator maps each question and ranked-gene profile to the most suitable specialized expert or expert combination in the available set.

#### Expert execution

The shared backbone is loaded once, and expert-specific domain-alignment and specialization adapters are activated on demand. Expert outputs are converted to a common textual schema containing the predicted answer, supporting genes, negative markers, tissue context, confidence, and execution metadata. The common schema enables heterogeneous experts to be composed without aligning their hidden representations.

### 4.7 Textual Optimization of the Playbook

The playbook *P* contains explicit natural-language rules for expert routing, tool construction, evidence synthesis, ambiguity handling, and abstention. We optimize this textual context directly while keeping the parameters of the orchestrating LLM fixed. This design follows recent work that treats prompts or textual context as optimizable objects using natural-language feedback rather than parameter gradients (Pryzant et al. 2023; Yang et al. 2024; Yuksekgonul et al. 2025; Zhang et al. 2025).

At optimization iteration *t*, example *i* is processed with the current playbook *P*_*t*_:

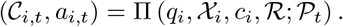

A textual-loss module then compares the output with the available training supervision:

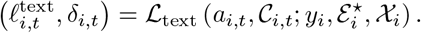

Here, 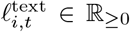 is a scalar error summary, *δ*_*i,t*_ is a structured natural-language diagnosis, and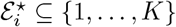 is the set of oracle experts used only by the training-time evaluator.

For an optimization mini-batch 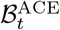, the diagnoses are distilled into recurrent, actionable insights:

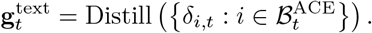

The operator Distill summarizes recurring diagnoses, and 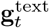 is the resulting textual gradient. It is not a numerical derivative; it is a natural-language surrogate that identifies how the current playbook should change to reduce routing, grounding, synthesis, or abstention errors.

A textual optimizer updates the playbook:

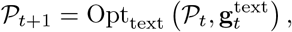

where Opt_text_ is the rule-editing operator. It can add, rewrite, merge, remove, or reorder rules while eliminating redundant and contradictory instructions. Detailed formalization of the textual loss, edit space, and discrete optimization interpretation is given in Appendix B.4.

Oracle experts, reference answers, textual losses, and optimizer feedback are unavailable at inference time. The final policy operates only on the user request, input profiles, expert descriptions, expert outputs, and the optimized playbook.

### 4.8 Extensibility

The framework is designed for expert-level extensibility: separately constructed specialists communicate through a shared textual interface. This modularity follows the principle of reusable and composable adapters (Houlsby et al. 2019; Pfeiffer et al. 2020, 2021). The present study evaluated a fixed set of experts; expansion to previously unseen experts remains future work.

## A Additional Expert-Specialization Results

Table 2 reports the tissue-level exact-match and grounding results for the expert-specialization benchmark. The table is placed in the appendix to keep the main Results focused on aggregate findings and cross-model comparisons.

**Table 2.**
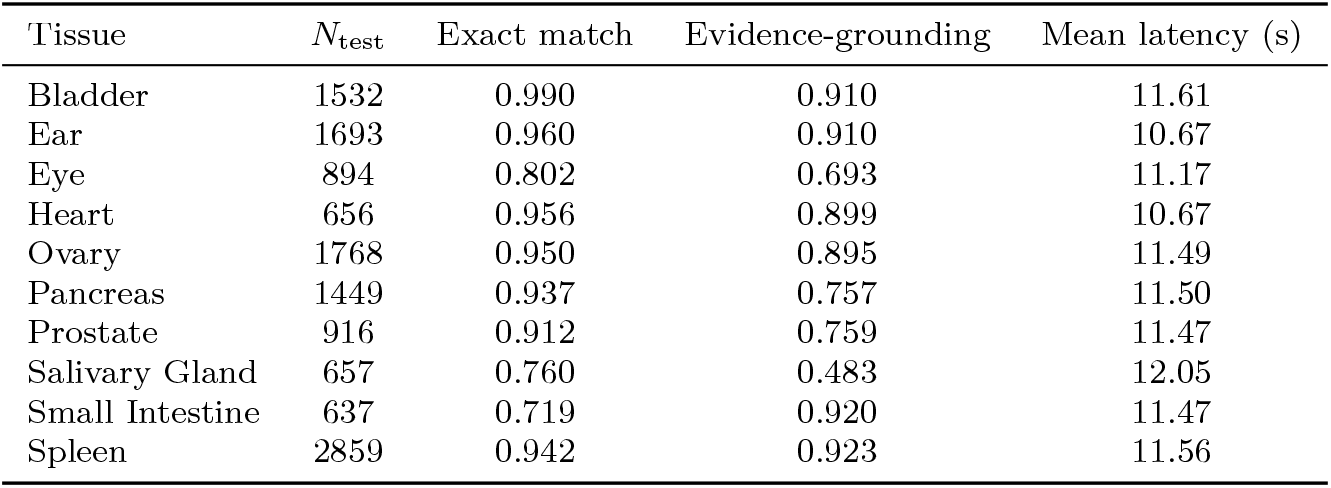
Stage 2 cell-type annotation performance across ten Tabula Sapiens tissues. Exact match is the primary task metric. Evidence-grounding reports the fraction of ground-truth evidence genes recovered by the generated rationale.

| Tissue | $N_{\text{test}}$ | Exact match | Evidence-grounding | Mean latency (s) |
| --- | --- | --- | --- | --- |
| Bladder | 1532 | 0.990 | 0.910 | 11.61 |
| Ear | 1693 | 0.960 | 0.910 | 10.67 |
| Eye | 894 | 0.802 | 0.693 | 11.17 |
| Heart | 656 | 0.956 | 0.899 | 10.67 |
| Ovary | 1768 | 0.950 | 0.895 | 11.49 |
| Pancreas | 1449 | 0.937 | 0.757 | 11.50 |
| Prostate | 916 | 0.912 | 0.759 | 11.47 |
| Salivary Gland | 657 | 0.760 | 0.483 | 12.05 |
| Small Intestine | 637 | 0.719 | 0.920 | 11.47 |
| Spleen | 2859 | 0.942 | 0.923 | 11.56 |

### A.1 Representation-Preservation Ablation

We compared expert specialization with and without the representation-preservation objective. The frozen domain-aligned representation achieved a held-out hidden-space Recall@10 of 0.969 before specialization. Without preservation regularization, Recall@10 decreased to 0.915. The regularized expert retained Recall@10 at 0.969 and achieved a lower validation loss than the unregularized model (1.143 versus 1.185; Table 3).

**Table 3.**
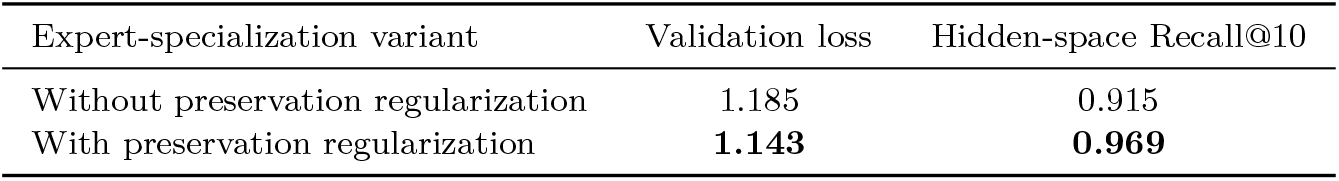
Representation-preservation ablation during expert specialization. Validation loss measures task fitting (lower is better), whereas hidden-space Recall@10 measures retention of the domain-aligned neighborhood structure (higher is better). The frozen domain-aligned representation had a pre-specialization Recall@10 of 0.969.

## B Technical Details of the Method

### B.1 Backbone and Prompt Representation

Let *s* be a tokenized prompt, let *L*(*s*) be its number of non-padding tokens, and let *d* be the hidden dimension of the causal language-model backbone. Given frozen backbone parameters *θ* and active adapter parameters Δ, the backbone produces

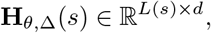

where row *r* contains the hidden state of prompt token *r*. We use the final non-padding hidden state as the prompt representation:

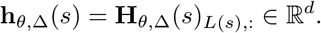

The notation

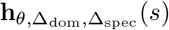

denotes the same backbone with the domain-alignment adapter Δ_dom_ followed by the specialization adapter Δ_spec_. The backbone parameters *θ* remain frozen throughout both adaptation components.

### B.2 Expanded Domain-Alignment Objective

For cell *i* and view *v* ∈ {1, 2}, define

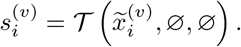

Let 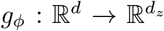 be the projection head, where *d*_*z*_ is the contrastive embedding dimension. The normalized projected representation is

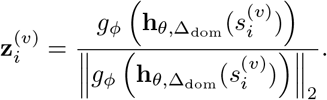

For a contrastive mini-batch of *N* cells, define the set of 2*N* augmented-view indices

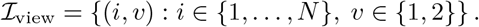

For view *v*, let 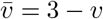 denote the other view of the same cell. Because all projected vectors are normalized, cosine similarity between vectors **u** and **v** is

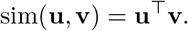

The domain-alignment loss is

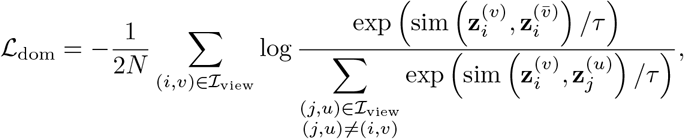

where *τ >* 0 is the contrastive temperature. The other view of the same cell is the positive sample, and every remaining view in the mini-batch is a negative sample.

### B.3 Expanded Expert-Specialization Objective

Let ℬ_spec_ be a specialization mini-batch and let *B* = _spec_ | ℬ | be its number of examples. For example *i*, the target completion is

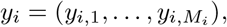

where *M*_*i*_ is its token length. The token-normalized completion loss is

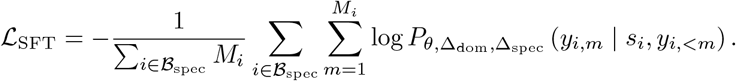

For each prompt *s*_*i*_, define the normalized current representation

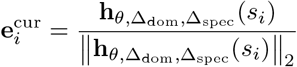

and the frozen domain-aligned reference

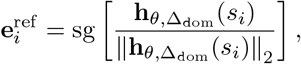

where sg[·] denotes stop-gradient.

#### B.3.1 Pointwise Alignment

The pointwise anchor loss is

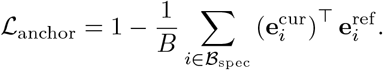

### B.3.2 Distribution Alignment

For state *σ* ∈ {cur, ref}, define the mini-batch mean

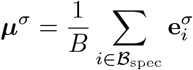

and componentwise variance

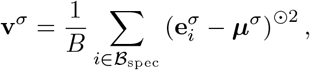

where (·)^*⊙*2^ denotes elementwise squaring. The distribution-alignment loss is

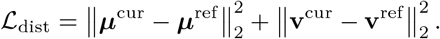

#### B.3.3 Relational Alignment

Choose any fixed ordering *i*_1_, …, *i*_*B*_ of the examples in ℬ_spec_. For *σ*∈ {cur, ref}, stack their normalized representations row-wise:

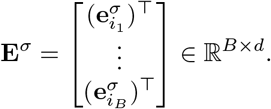

The pairwise cosine-similarity matrix is

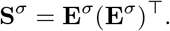

Let **1**_*B*_ ∈ ℝ^*B*^ be the all-ones vector, **I**_*B*_ ∈ ℝ^*B×B*^ the identity matrix, and

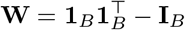

the off-diagonal mask. The relational loss is

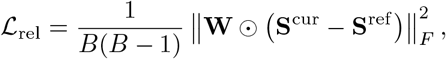

where ⊙ denotes elementwise multiplication and∥ · ∥ _*F*_ is the Frobenius norm. The diagonal is excluded because both similarity matrices have unit self-similarity.

The full specialization objective is

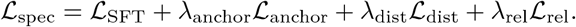

### B.4 Detailed Textual Optimization Formulation

Let

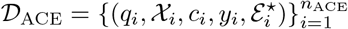

denote the orchestration optimization set, where *n*_ACE_ is the number of optimization examples and 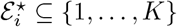 is the oracle expert set available only to the training-time evaluator.

At iteration *t* ∈ {0, …, *T*_opt_ − 1}, the textual policy produces

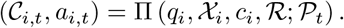

The textual-loss module returns

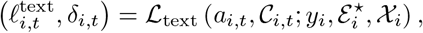

where 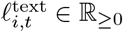 is the scalar loss and *δ*_*i,t*_ is its structured textual diagnosis.

For optimization mini-batch 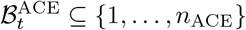, the diagnoses are distilled as

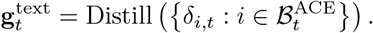

Although termed a textual gradient, 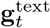 is not obtained by differentiation. It is a natural-language surrogate that summarizes a direction of improvement in the discrete playbook space.

The playbook is updated by

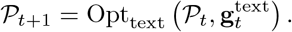

The allowed edit actions form the set

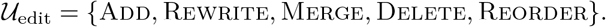

These actions alter routing rules, profile-specific constraints, evidence-synthesis instructions, and abstention criteria.

#### B.4.1 Discrete Optimization Interpretation

Let

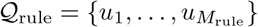

be an abstract collection of *M*_rule_ candidate textual rules, and let

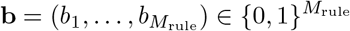

indicate which candidate rules are active. The playbook induced by **b** is

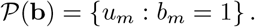

Under this abstraction, playbook selection can be written as

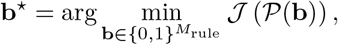

where the empirical evaluation objective is

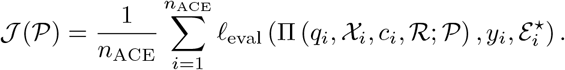

Here, *l*_eval_ is the scalar evaluator used to compare a policy output with its reference answer and oracle expert set.

The implemented optimizer operates over a richer space than binary rule selection because rules can also be rewritten, merged, or specialized. Let *N*(*P*_*t*_) denote the textual neighborhood induced by one or more actions in *U*_edit_, and let 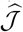 denote the textual optimizer’s local surrogate of *J*. Each update can be viewed as

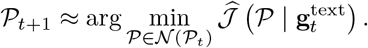

After *T*_opt_ optimization iterations, the final playbook is

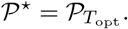

For an unseen query, inference is

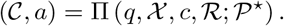

No reference answer, oracle expert identity, textual loss, or optimizer feedback is used during inference.

## C Metric Definitions

### C.1 Domain-Alignment Metrics

Let 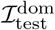 be the index set of held-out cells in the domain-alignment benchmark, and let *Y*_dom_ be its set of evaluated cell-type labels. Transfer predictions are obtained using a *k*_clf_-nearest-neighbor classifier, where *k*_clf_ denotes the fixed number of neighbors used by the classifier.

For class *c* ∈ *Y*_dom_, let TP_*c*_, FP_*c*_, and FN_*c*_ denote the numbers of true positives, false positives, and false negatives, respectively. Precision, recall, and F1 are

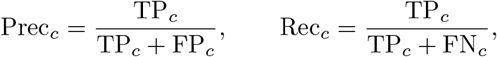

and

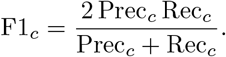

Macro-F1 is

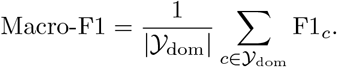

For Recall@5, let *N*_5_(*i*) be the set of five nearest training cells to held-out cell *i*, and let 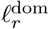 denote the reference cell-type label of any training or test cell indexed by *r*. We define

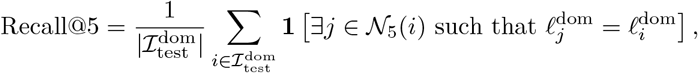

where **1**[·] is the indicator function.

### C.2 Expert-Specialization Metrics

Let 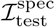 be the index set of held-out specialization examples. Let 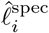 and 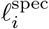 denote the generated and reference final labels for example *i*. Exact-match accuracy is

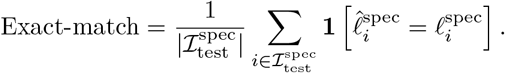

Let 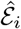 be the normalized, deduplicated set of genes returned in the generated evidence field, *G*_*i*_ the set of genes in the ranked input profile, and ℰ_*i*_ the reference evidence set. The per-example evidence-in-input score is

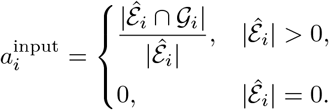

The dataset-level evidence-in-input rate is

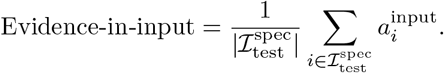

The per-example evidence-grounding score is

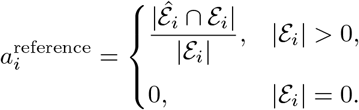

The dataset-level evidence-grounding rate is

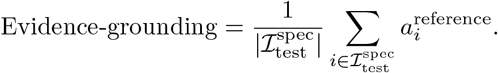

### C.3 Orchestration Metrics

Let 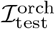 be the index set of held-out orchestration examples. The primary orchestration metrics quantify label compatibility, evidence agreement, marker coverage and the combined quality of the generated answer. Routing diagnostics are reported separately to isolate expert selection from answer synthesis.

#### C.3.1 Ontology-Aware Label Agreement

Let 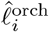 and 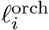 denote the predicted and reference cell-type labels for orchestration example *i*. Let *ν*(·) be the fixed label-normalization mapping used by the evaluator. Define

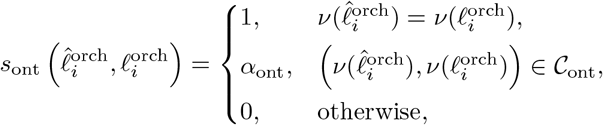

where *C*_ont_ is the fixed set of ontology-compatible label pairs and 0 *< α*_ont_ *<* 1 is the fixed partial-credit value. Hierarchical cell-type ontologies provide standardized relations for such comparisons (Diehl et al. 2016).

The dataset-level metric is

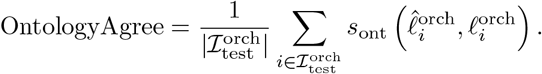

#### C.3.2 Input-Grounded Support-Gene Score

Let 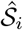 be the normalized, deduplicated set of support genes returned for orchestration example *i*, and let 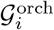 be the set of genes in the corresponding ranked input profile. The per-example score is

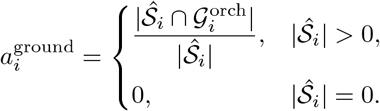

Assigning zero to an empty support set prevents a model from obtaining a perfect grounding score by omitting evidence entirely.

The dataset-level score is

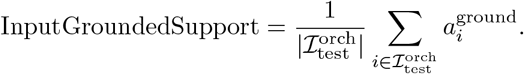

The complementary input-unsupported citation rate is

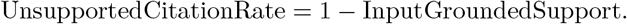

This metric evaluates whether cited genes are licensed by the observed input profile. It does not measure whether an input-present gene is biologically specific for the predicted label; biological label compatibility is evaluated separately through OntologyAgree.

#### C.3.3 Evidence Alignment

Evidence alignment measures whether the structured evidence returned by the model matches the reference support and exclusionary-marker fields. Let 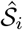 and *S*_*i*_ denote the predicted and reference support-gene sets for orchestration example *i*, and let 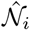 and *N*_*i*_ denote the corresponding predicted and reference negative-marker sets. For any two sets *A* and *B*, define

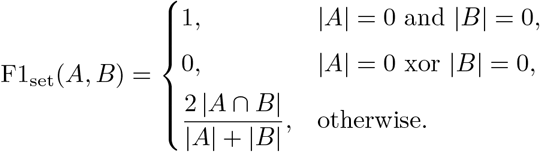

The support-gene and negative-marker alignment scores are

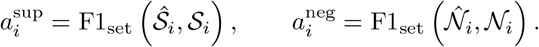

The per-example evidence-alignment score is

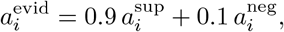

The 0.9/0.1 weighting prioritizes supporting genes because they provide direct positive evidence for the assigned identity, while retaining negative markers as secondary exclusionary evidence. The dataset-level metric is

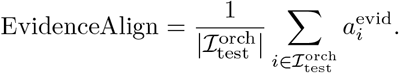

#### C.3.4 Marker Mention Coverage

Marker mention coverage measures whether the free-form rationale mentions the reference support and negative markers. Let *r*_*i*_ be the generated free-form answer. For a set of markers *M*, define

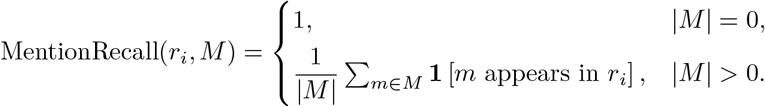

The per-example marker-coverage score is

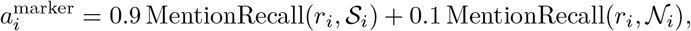

and the dataset-level score is

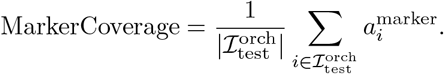

#### C.3.5 Expert Routing Quality

Let *C*_*i*_ be the set of experts actually called for example *i*, and let 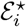 be the oracle expert set stored in the request metadata. If no oracle set is available, the evaluator uses the candidate expert set 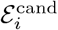. Define the target expert set

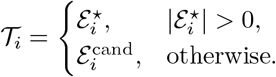

The routing recall and precision are

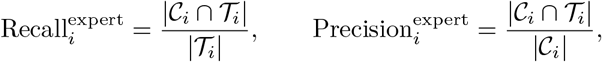

with the empty-set cases handled by assigning perfect routing only when no expert is required and no expert is called. Let 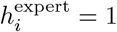 if at least one target expert is called and zero otherwise. The routing-quality score is

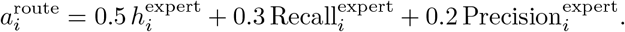

The reported routing quality is the mean over the held-out orchestration set,

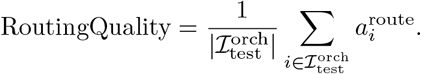

For routing-only controls, the same score is recomputed after replacing the observed expert-call trace with oracle, random or empty expert-call traces. These controls isolate expert selection and do not regenerate the final answer.

#### C.3.6 Nuanced Overall Score

The nuanced overall score is the weighted evaluator composite used to summarize the quality of each orchestration answer. Let 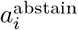 be the abstention-calibration score, 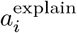 be the token-F1 similarity between the generated rationale and reference rationale, and 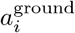 be the input-grounded support-gene score defined above. The per-example nuanced score is

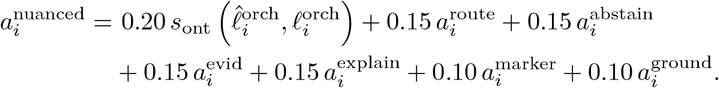

The reported nuanced overall score is the mean of 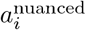 over 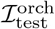.

#### C.3.7 Statistical Reporting

Pairwise model comparisons use paired randomization tests on the per-example metric differences. The reported tests compare scACORN with GPT-5 because GPT-5 is the strongest non-scACORN baseline in the main orchestration comparison.

#### C.3.8 Checkpoint Reporting

Pre-optimization values were computed using the initial playbook *P*_0_, and the final endpoint was specified before evaluation. Intermediate checkpoints were used to describe the optimization trajectory and were not used to select the reported playbook on the held-out benchmark.

## D Qualitative scACORN Interactions

We selected the following held-out Orchestration examples to illustrate the question-answering behavior supported by scACORN. To keep the appendix readable, we omit the complete ranked-gene lists and report only the user question, scACORN response, and reference answer.

### D.1 High-Agreement Examples

#### D.1.1 Single-Cell Endothelial Query

##### User question

I pasted a ranked gene list above and I am not sure what cell type this profile represents. Could you suggest the most likely cell identity and point to which markers in the list support that assignment? I am trying to decide between an endothelial versus a mesenchymal or smooth-muscle interpretation.

##### BioAgent

Most likely identity: cardiac endothelial cell (predicted tissue: heart). This assignment is supported by endothelial markers present in the ranked list, including VWF, ENG, and EMCN. Mesenchymal and smooth-muscle alternatives are weakened by the absence of COL1A1, DCN, ACTA2, and TAGLN from the ranked profile. Confidence: high. Plausible alternative labels include generic endothelial cell, vascular endothelial cell, capillary endothelial cell, and vein endothelial cell.

##### Ground truth

Most likely cell identity: cardiac endothelial cell. This assignment is supported by VWF and EMCN being present in the ranked genes, with ENG further reinforcing an endothelial signature. The lack of COL1A1 provides little support for a fibroblast interpretation.

#### D.1.2 Multi-Cell Immune Versus Myeloid Query

##### User question

I pasted two ranked gene lists above corresponding to profile IDs A and B. For each profile ID, provide a single best cell-type annotation with a confidence level and short rationale, and explicitly list the supporting evidence and negative markers used from each list.

##### BioAgent

Profile A—naive thymus-derived CD4-positive, alpha-beta T cell (confidence: high). The ranked list contains IL7R and LTB, which support a naive CD4 T-cell interpretation, while effector and activation alternatives are weakened by the absence of GZMB, IFNG, IL4, and IL5.

Profile B—mononuclear phagocyte (confidence: moderate). Antigen presentation markers including CD74, HLA-DRA, HLA-DRB1, and CST3 support an antigen-presenting mononuclear phagocyte state. B-cell and activated T-cell alternatives are weakened by the absence of canonical lymphoid and cytotoxic markers.

##### Ground truth

Profile A: naive thymus-derived CD4-positive, alpha-beta T cell—high confidence. IL7R and LTB are present, while canonical effector or activation markers including IFNG, IL4, IL5, and GZMB are not detected among the top-ranked genes.

Profile B: mononuclear phagocyte—high confidence. AIF1, TYROBP, and FCER1G appear in the ranked list, while lymphoid and endothelial markers including CD3D, MS4A1, PECAM1, and EPCAM are absent or low.

### D.2 Biologically Distinctive Examples

#### D.2.1 Single-Cell Retinal Query

##### User question

I pasted a ranked gene list from a cluster above. Can you give the most likely cell identity and one or two plausible alternative annotations, with a brief explanation of the evidence?

##### BioAgent

Most likely identity: eye photoreceptor cell (predicted tissue: eye). This assignment is supported by neuronal genes SNAP25 and MAP2 in the ranked list. S100B, TMEM119, and COL1A1 are absent, weakening glial and fibroblast interpretations. Plausible alternatives include a generic retinal neuron or retinal bipolar neuron. Confidence: moderate because the detected neuronal genes are not fully photoreceptor specific.

##### Ground truth

Most likely identity: eye photoreceptor cell. MAP2 and SNAP25 support a neuronal retinal identity. The absence of glial or stromal markers such as S100B and COL1A1 argues against a glial or fibroblast interpretation. Plausible alternatives include other retinal neurons such as bipolar or amacrine cells because MAP2 and SNAP25 are neuronal but not fully photoreceptor specific.

#### D.2.2 Multi-Cell Epithelial Versus Perivascular Query

##### User question

I pasted two ranked gene lists above for profiles A and B. Could you annotate each profile and determine whether they represent related states within one lineage or clearly distinct cell identities?

##### BioAgent

Profile A—enterocyte of the epithelium proper of the ileum. The ranked genes include APOA1, APOA4, SLC5A1, FABP2, and ANPEP, supporting an absorptive ileal enterocyte identity. Confidence: high.

Profile B—pericyte. The ranked genes include RGS5, NOTCH3, SPARC, IGFBP7, and CALD1, supporting a perivascular mesenchymal identity. Confidence: moderate.

The two profiles represent clearly distinct cell identities rather than related states within one lineage.

##### Ground truth

Profile A: enterocyte of the epithelium proper of the ileum. APOA1, APOA4, and SLC5A1 support an absorptive enterocyte identity, whereas MUC2 and CHGA are not enriched and argue against goblet or enteroendocrine interpretations.

Profile B: pericyte. RGS5, NOTCH3, and COL4A1 support a pericyte or perivascular program, whereas PECAM1 and VWF are not enriched and argue against an endothelial identity.

The two profiles are clearly distinct cell identities rather than related states of one lineage.

## Code availability

The code is available at https://github.com/arashrasti96/scACORN.

